# Robust Cooperative Learning for Multiview Regression via Soft-IPOD

**DOI:** 10.64898/2026.09.24.754279

**Authors:** Shunsuke Yoshimura, Tomoyuki Hiroyasu, Kensuke Tanioka

**Affiliations:** Graduate School of Life and Medical Sciences, Doshisha University, Kyoto, Japan; Department of Biomedical Sciences and Informatics, Doshisha University, Kyoto, Japan

## Abstract

Multiview data analysis has become increasingly important for integrating heterogeneous information from multiple data sources. Cooperative learning provides a flexible framework for multiview integration by bridging early and late fusion strategies. However, its use of squared loss makes it sensitive to outliers. In this study, we propose robust cooperative learning based on the soft-IPOD framework. The proposed method introduces an observation-specific outlier absorption vector into the cooperative learning objective, thereby reducing the influence of outlying observations while preserving information sharing across views. The resulting optimization problem can be formulated as a weighted Lasso-type problem using an augmented design matrix and can be naturally extended to more than two data views. Numerical simulations and a real-data application to labor onset prediction demonstrate the predictive performance of the proposed method in the presence of outliers.

## 1 Introduction

Recent years have witnessed growing interest in multiview data analysis in statistics and machine learning, where data obtained from multiple distinct information sources for the same subjects are jointly integrated and analyzed [1, 2, 3, 4]. For example, in biomedical and life science research, it has become increasingly common to collect information from different modalities, including genomics, transcriptomics, proteomics, and imaging data, for the same subjects [2, 3, 4]. By appropriately integrating multiple data views, complementary and shared information that cannot be captured by a single view alone can be exploited, potentially improving predictive performance for an outcome of interest [1, 2, 4]. However, determining when and how to integrate information across multiple views remains a fundamental challenge in multiview data analysis [1].

Representative data integration strategies in supervised learning include early fusion and late fusion [1]. In early fusion, features from multiple views are combined into a single feature space prior to model fitting, and a unified predictive model is then constructed using the integrated data [2, 5, 3, 4]. Alternatively, view-specific features can first be transformed into lower-dimensional representations before integration, using methods such as principal component analysis or autoencoders [6, 7]. This approach is relatively straightforward to implement and allows information from different views to be utilized simultaneously. However, in high-dimensional settings, it may suffer from an increased number of features and the accumulation of noise. In contrast, late fusion constructs a separate predictive model for each view and subsequently combines the predictions obtained from these individual models [8, 9, 10, 11]. Various extensions of late fusion have also been considered, including stacking-based approaches [12]. This strategy offers the flexibility to employ different modeling approaches for different views; however, it may fail to fully exploit information shared across views during model construction.

To provide a unified framework for these two approaches, Ding et al. [1] proposed cooperative learning. Cooperative learning introduces an agreement penalty into the objective function, in addition to the usual prediction loss, to encourage agreement between the predictions obtained from different views. This allows predictive models to be constructed by leveraging information across views while maintaining predictive accuracy for the outcome. By adjusting the strength of the agreement penalty, cooperative learning provides a continuum between early fusion and late fusion. Thus, cooperative learning provides a flexible framework for controlling the degree of information sharing across views, thereby bridging early fusion and late fusion. Related ideas that exploit information sharing or relationships across views have also been considered in other multiview methods, including collaborative regression [13], co-regularization [14], and sparse integrative discriminant analysis (SIDA) [15].

However, in practical data analysis, the error distribution may deviate from normality, and the observed data may contain outliers. In such situations, estimation based on the squared-error loss can be substantially affected by a small number of outlying observations because large residuals receive disproportionately large weights. To address this issue, various robust methods have been proposed to reduce the influence of outliers and departures from the assumed error distribution. Representative approaches include robust estimation based on the Huber loss[16, 17], quantile regression[18, 19], regularized least absolute deviation methods such as LAD-Lasso[20], and divergence-based robust estimation[21, 22]. Another approach is the penalized regression framework proposed by She and Owen [23], which introduces observation-specific parameters representing outlier effects and imposes a sparsity-inducing penalty on these parameters, thereby enabling outlier detection and robust regression simultaneously. A representative method within this framework is soft-IPOD. This framework has also been extended to sparse regression settings, where sparsity is imposed on both the regression coefficients and the outlier effects [24].

In this study, we focus on the soft-IPOD approach. Soft-IPOD achieves robustness by augmenting a standard regression model with observation-specific parameters that absorb the effects of outlying observations and imposing a penalty on these parameters. This formulation can be naturally incorporated into the objective function of cooperative learning without substantially altering its original framework. Consequently, the cooperative learning framework can be extended to provide robust multiview prediction while retaining its connection to both early fusion and late fusion. Based on this idea, we propose a robust extension of cooperative learning, referred to as robust cooperative learning.

The contributions of this paper are summarized as follows. First, we propose a robust extension of cooperative learning that incorporates an outlier absorption mechanism into the regression model. Second, we develop an efficient optimization procedure based on an augmented design matrix, which allows the proposed model to be solved as a weighted Lasso-type problem. Third, we extend the proposed framework to settings with more than two data views and provide a unified formulation. Finally, we demonstrate the effectiveness of the proposed method through numerical simulations and a real-data application to labor onset prediction.

The remainder of this paper is organized as follows. Section 2 introduces the proposed robust cooperative regularized linear regression model for two data views, including its formulation and optimization procedure, and discusses its relationship with early and late fusion strategies. Section 3 extends the proposed framework to more than two data views and presents the corresponding optimization procedure. Section 4 presents the numerical simulation study, including the simulation design and results. Section 5 applies the proposed method to a real dataset on labor onset prediction and compares its performance with baseline methods. Finally, Section 6 provides a discussion of the results and concludes the paper.

## 2 Robust cooperative learning with two data views based on soft-IPOD

In this section, we show the framework and the algorithm of robust cooperative learning based on two data block matrices. We first present the formulation of the regularized linear regression model and the corresponding optimization problem. We therefore describe the estimation procedure for the model parameters and provide the detailed algorithms for solving optimization problems. Next, we explain the relationship between the hyperparameter *ρ* and the early fusion and late fusion strategies in the same manner as cooperative learning.

### 2.1 Robust cooperative regularized linear regression

In this subsection, we describe the formulation of the proposed robust cooperative regularized linear regression model. Let 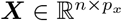and 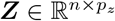 denote two feature matrices (data views), and let ***y*** ∈ ℝ^*n*^ denote the response vector. We consider the problem of jointly estimating the regression coefficient vectors 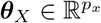 and 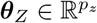, together with an outlier absorption vector ***γ*** ∈ R^*n*^. We assume that the columns of ***X*** and ***Z*** are standardized and that the response vector ***y*** is centered so that it has mean zero. Under this assumption, the intercept term can be omitted without loss of generality. Furthermore, ∥ · ∥_2_ denotes the Euclidean (*ℓ*_2_) norm defined for a vector ***v*** by 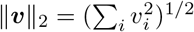 and ∥ · ∥_1_ denotes the *ℓ*_1_ norm defined by ∥ ν ∥_1_ = Σ_i_ |ν_i_ |. The parameters are obtained by minimizing the following objective function:

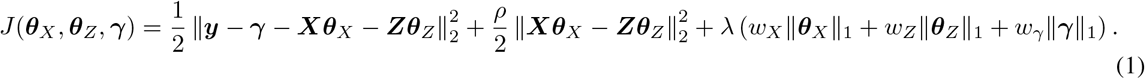

In Equation (1), the first term represents the squared prediction error after accounting for the sample-wise outlier effects represented by γ, while the second term is an agreement penalty that encourages the view-specific predictions ***Xθ***_*X*_ and ***Zθ***_*Z*_ to be similar. The last term is a weighted *ℓ*_1_ regularization penalty. Here, *λ* controls the overall strength of regularization, whereas *w*_*X*_, *w*_*Z*_, and *w*_*γ*_ are penalty factors. These penalties induce sparsity in the regression coefficients and in the outlier vector. The vector ***γ*** captures deviations from the Gaussian noise assumption and absorbs potential outliers in the response variable.

However, the squared loss is sensitive to large deviations, and even a small number of extreme observations may distort the estimation of regression coefficients[23]. By introducing the mean-shift vector ***γ*** with an *ℓ*_1_ penalty, the proposed model allows only a limited number of observations to deviate from the main regression structure. Consequently, potential outliers are absorbed by ***γ***, while the cooperative regression coefficients ***θ***_*X*_ and ***θ***_*Z*_ are estimated based on the main data structure. Importantly, incorporating the Soft-IPOD formulation does not alter the essential structure of the cooperative learning objective, but only augments it through the addition of the mean-shift term[23]. Therefore, the proposed robust extension preserves the original fusion continuum controlled by the agreement parameter *ρ*. In particular, the framework continues to span the spectrum between early fusion and late fusion as special cases, while providing robustness against outlying observations in the response variable. This property ensures that the interpretability and flexibility of cooperative learning are maintained while improving the stability of the estimation under outliers. The hyperparameter *ρ* ≥ 0 controls the strength of agreement between the predictions obtained from the two data views. When *ρ* = 0, the model reduces to early fusion, whereas *ρ* = 1 corresponds to late fusion.[1] Intermediate values of *ρ* interpolate between these two regimes. Thus, the proposed model provides a flexible framework that interpolates between early fusion and late fusion. A more detailed discussion of these regimes will be provided in Section2.2.

Equation (1) defines a convex optimization problem when *ρ* ≥ 0, *λ* ≥ 0, and all penalty factors are nonnegative. For efficient computation, we rewrite Equation (1) using an augmented design matrix as follows:

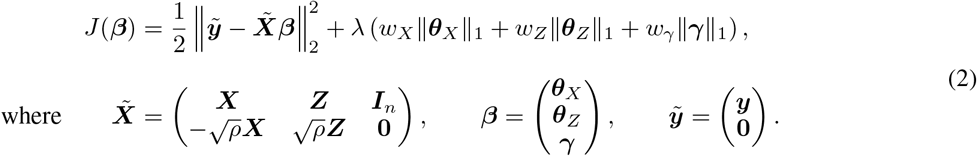

This augmented representation shows that the proposed optimization problem can be formulated as a weighted Lasso-type penalized regression problem with an augmented design matrix. For each candidate combination of the agreement parameter, the global regularization parameter, and the penalty factors, the model parameters are estimated using standard algorithms for *ℓ*_1_-penalized regression. The tuning parameters are then selected by cross-validation [25, 26]. Algorithm 1 summarizes the direct algorithm based on this augmented formulation, including the cross-validation procedure. In this algorithm, we focus on the gamma-PF setting, where the penalty factors for the regression coefficients are fixed as *w*_*X*_ = *w*_*Z*_ = 1, while the penalty factor for the outlier vector, *w*_*γ*_, is selected through an outer grid search.

#### Algorithm 1

Direct algorithm for robust cooperative regularized regression with γ-penalty factors

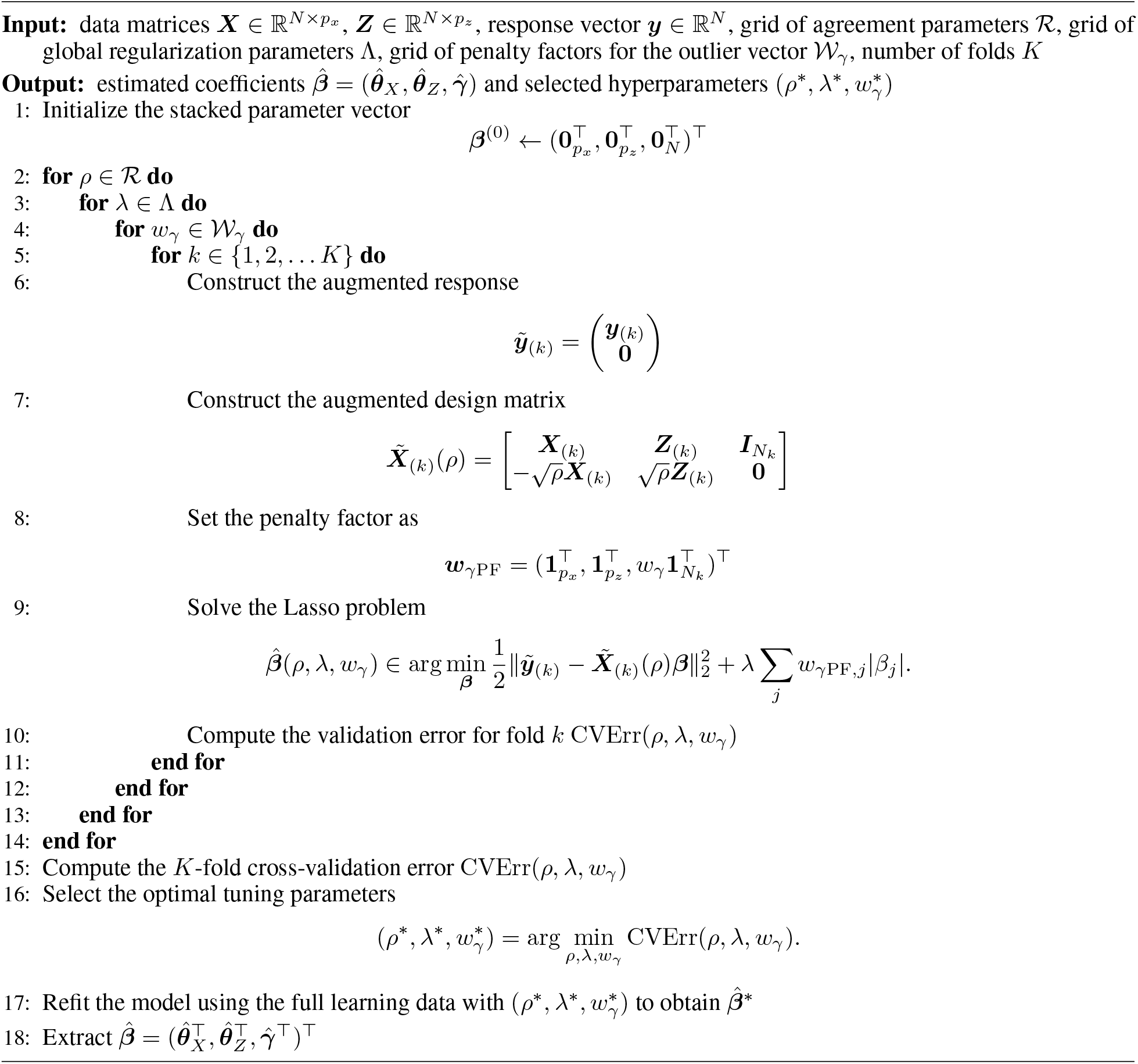

For each candidate value of the agreement parameter *ρ*, the global regularization parameter *λ*, and the penalty factor for the outlier vector *w*_*γ*_, an augmented design matrix and response vector are constructed. In this setting, the penalty factors for the regression coefficients are fixed as *w*_*X*_ = *w*_*Z*_ = 1, whereas *w*_*γ*_ is selected from an outer grid. The resulting optimization problem can then be written as a weighted Lasso-type regression problem with an augmented design matrix, which can be efficiently solved using coordinate descent algorithms implemented in standard software packages such as `glmnet` [26]. For each combination of (*ρ, λ, w*_*γ*_), the prediction error is evaluated using *K*-fold cross-validation on the learning data. The optimal tuning parameters (*ρ*^*∗*^, *λ*^*∗*^, 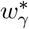) are selected as those minimizing the cross-validation error. Finally, the model is refitted using the full learning data with the selected tuning parameters.

### 2.2. Relation to early/late fusion

To clarify the relationship with existing fusion strategies, we temporarily ignore the regularization terms in Equation (1) and consider the objective function conditional on a fixed mean-shift vector ***γ***. Letting ***y***^*∗*^ = ***y***− γ, the proposed model can then be viewed as a cooperative learning model applied to the adjusted response ***y***^*∗*^. Consequently, conditional on a fixed γ, the proposed robust extension retains the relationship between the agreement parameter *ρ* and the early- and late-fusion regimes of the original cooperative learning framework [1]. For a fixed γ, define the augmented design matrix for the regression coefficient blocks as

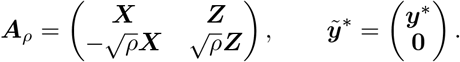

Then, ignoring the regularization terms, the objective function can be written as

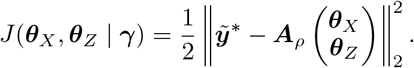

When *ρ* = 0, the agreement penalty vanishes and the objective reduces to a least-squares loss using the concatenated feature matrices.

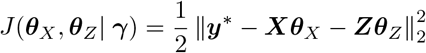

In this case, the prediction is obtained from a single model that uses all features simultaneously, which corresponds to the early fusion approach. Next, consider the case *ρ* = 1. From the definition of **A**_*ρ*_, we obtain

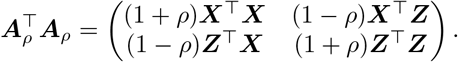

Assuming ***X*** and ***Z*** have full rank, and omitting the *ℓ*_1_ penalties, we obtain the least squares estimates

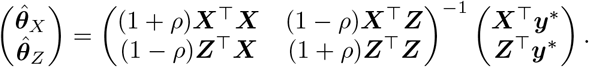

In the unregularized case with a fixed outlier vector γ, substituting *ρ* = 1 gives

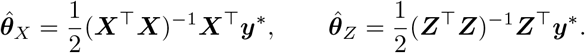

Substituting these expressions into the prediction rule 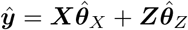 gives

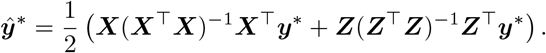

Let 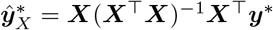 and 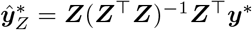 denote the predictions obtained by fitting separate least squares models using ***X*** and ***Z***, respectively. Then the final prediction becomes

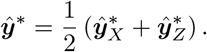

Thus, when *ρ* = 1, the proposed model produces the average of the predictions obtained from the two separate models, which corresponds to a late fusion strategy[1].

Therefore, the proposed framework provides a flexible mechanism that continuously bridges early fusion and late fusion. By selecting an appropriate value of *ρ*, the model can balance the benefits of joint modeling and view-specific learning while incorporating shared information between the data views.

## 3 Robust cooperative learning with more than two data views

In this section, we extend the robust cooperative regularized linear regression model to the case of multiple data views. This framework can be naturally extended to handle multiple data views while retaining robustness. Let 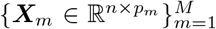 denote feature matrices corresponding to *M* data views, and let ***y*** ∈ ℝ^*n*^ denote the response vector. We estimate regression coefficient vectors 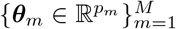 together with an outlier absorption vector γ ∈ ℝ^*n*^.

Assuming that each feature matrix is column-wise standardized and the response vector is centered, the intercept term can be omitted. For a fixed value of *ρ* ≥ 0 and *λ ≥* 0 the robust cooperative learning objective function for multiple data views is given by

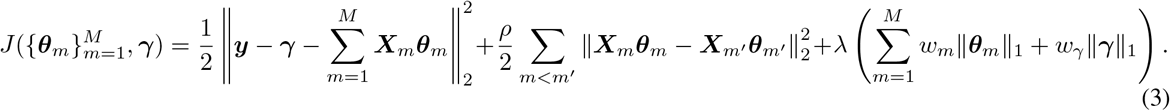

where *w*_*m*_ denotes the penalty factor for the regression coefficients of the *m*-th data view, and *w*_*γ*_ denotes the penalty factor for the outlier absorption vector.

The objective function in equation (3) is convex, and can be reformulated as a Lasso-type problem using an augmented design matrix. Let

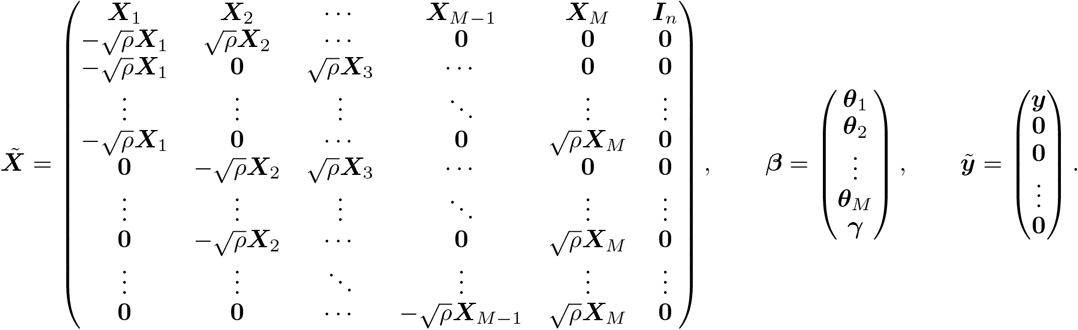

In this augmented design matrix, the first block row corresponds to the prediction error, while each subsequent block row corresponds to the agreement penalty between a distinct pair of data views. For each pair of views (*m, m*^*′*^) with 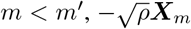 and 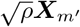^*′*^ are placed in the corresponding column blocks, respectively, while all other column blocks are set to zero matrices. Using these augmented quantities, the objective function in equation (3) can be rewritten as

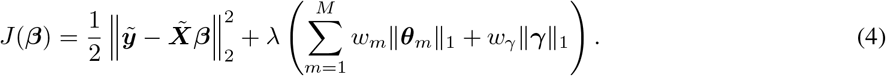

The augmented formulation can be solved directly as a weighted Lasso problem.

## 4 Numerical Simulation

We conducted numerical simulations to evaluate the predictive performance and robustness of the proposed method in high-dimensional multi-view settings. The simulation design was based on a latent factor model similar to that used in the cooperative learning literature, where multiple data views share a common latent structure. Furthermore, following the experimental setup of cooperative learning, we included both the original methods and their robust extensions in order to assess the impact of robustness under identical conditions.

### 4.1 Simulation design

#### 4.1.1 Data generation

For each simulation replicate, we generated *N* = 10000 observations and randomly selected *n*_train_ = 200 observations as the training set, with the remaining *n*_test_ = 9800 observations used as the test set. The relatively large test set was used to obtain a stable estimate of prediction error, while model fitting and tuning were performed using only the training data. The dimensions of the two data views were set to *p*_*X*_ = *p*_*Z*_ = 1000. Let 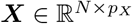and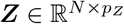denote the feature matrices for the two data views. We first generated noise matrices 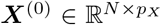 and 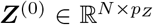, whose entries were independently sampled from a standard normal distribution. To introduce a shared latent structure between the two data views, the number of latent factors was set to *p*_*U*_ = 30. For *m* = 1, …, *p*_*U*_, the latent factor vectors were generated independently as

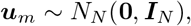

and the latent factor matrix was defined as 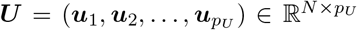.The generated latent factors were incorporated into the first 30 columns of ***X*** and ***Z*** (***x***_*·m*_ and ***z***_*·m*_), according to

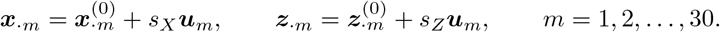

Here, *s*_*X*_ and *s*_*Z*_ are factor-loading parameters that control the strength of the latent signal contained in ***X*** and ***Z***, respectively. Larger values of *s*_*X*_ or *s*_*Z*_ correspond to a stronger contribution of the shared latent factors to the corresponding data view. No latent factors were added to columns 31 through 1000, which remained independent standard normal noise variables. Thus, among the 1000 variables in each data view, the first 30 contained latent structure associated with the response, whereas the remaining 970 were noise variables.

The true conditional mean was generated from the latent factors as

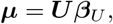

where ***β***_*U*_ = (8, …, 8)^*T*^ ∈ ℝ^30^. Consequently, the true signal was given by

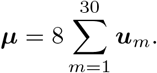

Before introducing outliers, the response variable was generated as

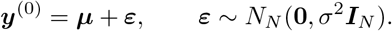

The signal-to-noise ratio (SNR) was defined as

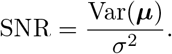

For each simulation replicate,

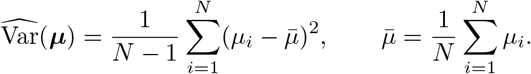

For a given value of SNR, the noise standard deviation was set to

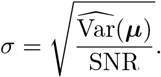

To evaluate robustness against outliers, contamination was introduced only into the responses in the training set. Let *π*_out_ denote the outlier contamination proportion. We randomly selected without replacement *n*_out_ = ⌊*π*_out_*n*_train_⌋ observations from the training set, where ⌊*a*⌋denotes the greatest integer less than or equal to *a*. For each selected observation *i*, a mean-shift perturbation was independently generated as

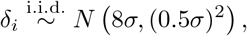

and the contaminated response was defined as

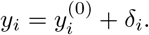

No outliers were introduced into the test responses, so that predictive performance could be evaluated on an uncontaminated test set.

Finally, the variables in ***X*** and ***Z*** were standardized using the means and standard deviations calculated from the training data, and the same scaling was applied to the test data.

#### 4.1.2 Simulation settings

We investigated the effects of the signal structure across the two data views, the signal-to-noise ratio (SNR), and the outlier contamination proportion on the performance of the competing methods. To vary the strength of the latent signal contained in the two data views, we considered three configurations of the factor-loading parameters:

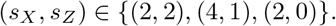

The setting (*s*_*X*_, *s*_*Z*_) = (2, 2) represents a balanced setting in which both data views contain latent signals of comparable strength. The setting (4, 1) represents an asymmetric setting in which both views contain latent signals, but the signal in ***X*** is stronger than that in ***Z***. Finally, the setting (2, 0) represents a setting in which only ***X*** contains the latent factors used to generate the response, whereas ***Z*** does not contain these response-generating latent factors. This simulation framework, in which the correlation and signal strength across data views are controlled through shared latent factors, follows the simulation design of the original cooperative learning study.

We considered three SNR values,

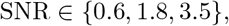

and three outlier contamination proportions in the training data,

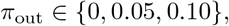

corresponding to contamination levels of 0%, 5%, and 10%, respectively. All combinations of the three factor-loading configurations, three SNR values, and three outlier contamination proportions were considered, resulting in a total of 3 × 3 × 3 = 27 simulation settings.

The hyperparameters for each method were selected by five-fold cross-validation using only the training data. For each simulation setting, 100 independent simulation replicates were performed. Predictive performance was evaluated using the mean squared error (MSE) with respect to the true signal ***µ*** in the test data.

We also evaluated the number of features selected by each method. In each simulation replicate, a feature was regarded as selected if its estimated regression coefficient was nonzero, and the total number of selected features was recorded. The distribution of the number of selected features across the 100 simulation replicates was then examined.

The selected agreement parameter *ρ* for cooperative learning was also examined as a supplementary analysis. For each candidate value of *ρ*, the selection proportion was calculated as the number of simulation replicates in which that value was selected by cross-validation divided by the total number of simulation replicates.

For the MSE and the number of selected features, summary statistics were calculated across the 100 simulation replicates. Detailed results for the number of selected features and the selected values of *ρ* are provided in Appendices A.2 and A.3, respectively.

#### 4.1.3 Comparison methods

To evaluate the predictive performance and robustness of the proposed method, we compared a total of 11 methods, consisting of six non-robust methods and five robust methods.

For the non-robust methods, we considered **Separate X** and **Separate Z**, in which Lasso regression was applied separately to each data view. We also considered **Early Fusion**, in which the features from the two data views were concatenated before fitting a Lasso model, and **Late Fusion**, in which separate prediction models were fitted to the two data views and their predictions were subsequently combined. In addition, we considered two versions of cooperative learning: **Cooperative Learning (no-PF)**, which uses common penalty factors for the regression coefficients, and **Adaptive Cooperative Learning (ad-PF)**, which uses adaptive penalty factors.

For the robust methods, we considered **Robust Separate X**(**Robust Sep X**), **Robust Separate Z**(**Robust Sep Z**), **Robust Early Fusion**(**Robust EF**), and **Robust Late Fusion**(**Robust LF**), which incorporate an outlier absorption term into the corresponding non-robust methods. We further considered the proposed **Robust Cooperative Learning** method. For Robust Cooperative Learning, the penalty factors for the regression coefficients were fixed at *w*_*X*_ = *w*_*Z*_ = 1, while only the penalty factor *w*_*γ*_ for the outlier absorption vector ***γ*** was tuned. Thus, an adaptive penalty-factor version of Robust Cooperative Learning was not considered in the present simulation study. In the figures, the proposed method under this setting is denoted by **Robust coop noPF**.

Predictive performance was compared among all 11 methods using the same simulated datasets. For clarity of presentation, the main text presents the MSE results for six methods: Early Fusion, Late Fusion, Cooperative Learning (no-PF), Robust Early Fusion, Robust Late Fusion, and Robust Cooperative Learning (Robust coop noPF). The MSE results for all 11 methods are provided in Appendix A.1.

For the variable-selection analysis, seven methods were compared, excluding Separate X, Separate Z, Robust Separate X, and Robust Separate Z, and the results are provided in Appendix A.2. In addition, for Cooperative Learning (no-PF), Adaptive Cooperative Learning (ad-PF), and Robust Cooperative Learning (Robust coop noPF), the distributions of the selected agreement parameter *ρ* are reported in Appendix A.3.

### 4.2 Simulation results

Figure 1 shows the test MSE for the six selected methods across all 27 simulation settings. Overall, the test MSE tended to decrease as the SNR increased. As the outlier contamination level increased, the test MSE of the non-robust methods generally increased, whereas the increase was relatively small for the robust methods.

**Figure 1:**
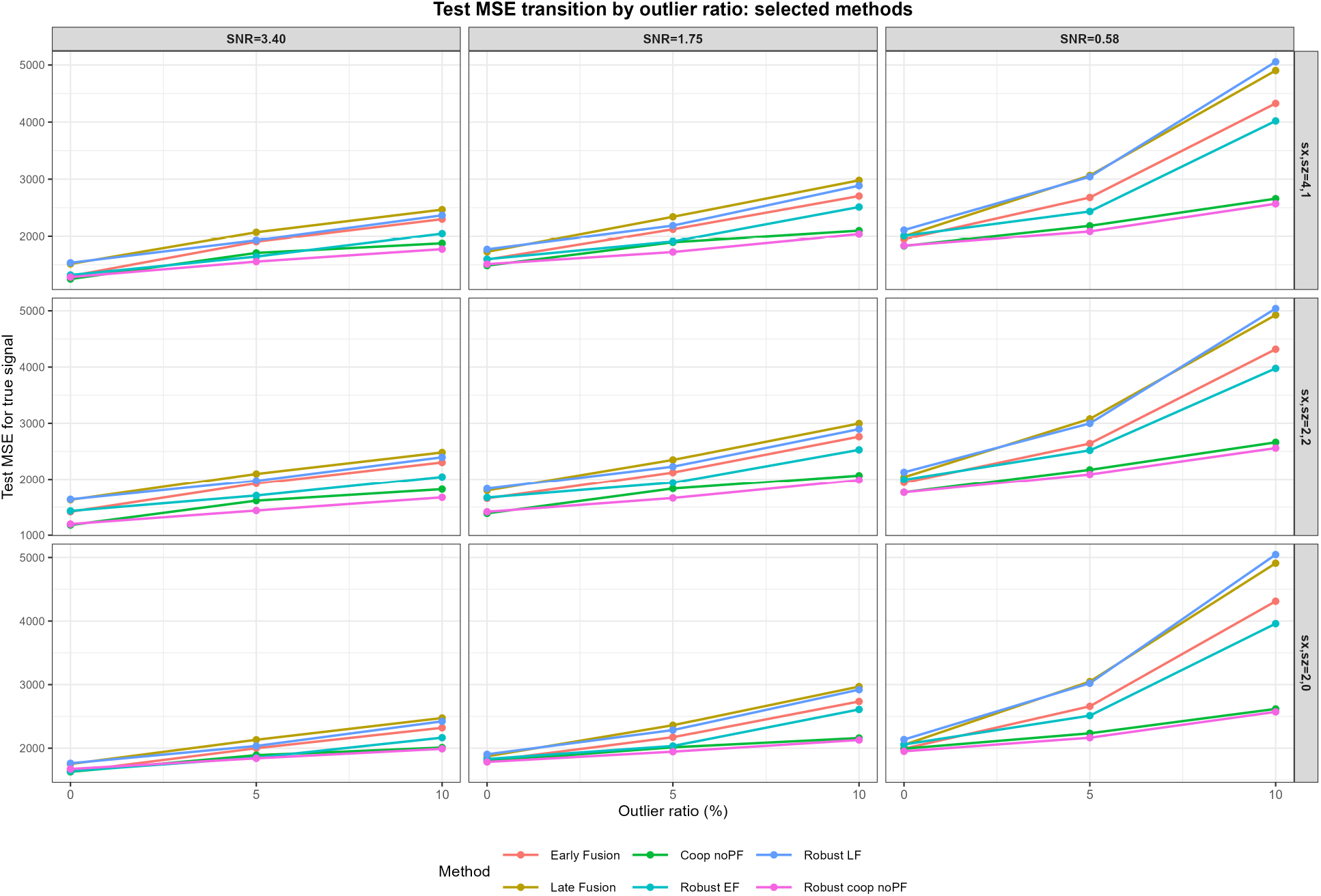
MSE transition for selected methods.

The relative performance of the methods also varied according to the signal structure across the two data views. When both views contained informative shared signals (*s*_*X*_ = 2, *s*_*Z*_ = 2), the proposed Robust Cooperative Learning method showed favorable predictive performance, particularly in the presence of outliers. A similar pattern was observed in the asymmetric setting (*s*_*X*_ = 4, *s*_*Z*_ = 1), where the two views differed in signal strength.

When the second view contained no shared latent signal (*s*_*X*_ = 2, *s*_*Z*_ = 0), the proposed method also maintained favorable predictive performance and achieved a lower test MSE than the separate-view methods in several settings.

Overall, the test MSE exhibited different patterns depending on the SNR, the outlier contamination level, and the signal structure across the data views. The results for the number of selected features and the selected agreement parameter *ρ* are provided in Appendices A.2 and A.3, respectively.

## 5 Real Data Application

### 5.1 Application to labor onset

This dataset was collected to predict the time to spontaneous delivery based on multi-omics measurements and is the same dataset as that used in the previous study on cooperative learning [27, 1]. It consists of longitudinal blood samples collected during the third trimester over a 120-day period from a cohort of 53 pregnant women who underwent spontaneous delivery.

The blood samples include 1,322 proteomic features and 3,529 metabolomic features, measured using non-targeted mass spectrometry, aptamer-based proteomics assays, and single-cell mass cytometry [27]. We defined the response variable as the number of days from sampling to the onset of spontaneous delivery and the explanatory variables as the proteomic and metabolomic measurements, forming a two-view dataset.

The dataset was randomly divided into a training set of 40 subjects and a test set of 13 subjects. Prior to analysis, following Ding et al. [1], the proteomic and metabolomic features were screened based on their variance across subjects.

The same 11 methods considered in the numerical simulations were used for comparison. For each data split, hyperparameters were selected by 10-fold cross-validation using only the training data, and predictive performance was evaluated on the test data using mean squared error (MSE). In addition, the number of features selected by each method was examined as an exploratory analysis.

The entire procedure, from data splitting to model evaluation, was repeated 10 times, and the MSE and the number of selected features were summarized across the 10 repetitions.

### 5.2 Comparison between the proposed method and baseline method

The results are shown in Table 1. Among all methods, the proposed Robust Cooperative Learning method achieved the lowest test MSE, indicating favorable predictive performance.

**Table 1:** Test MSE and the number of selected features (mean ± SD) on the labor onset dataset.

| Method | Test MSE (mean ± SD) | Selected features (mean ± SD) |
| --- | --- | --- |
| Separate X | 392.6 ± 283.2 | 18.8 ± 12.6 |
| Separate Z | 360.0 ± 170.3 | 18.1 ± 16.3 |
| Early Fusion | 662.4 ± 735.7 | 26.2 ± 16.0 |
| Late Fusion | 399.7 ± 231.2 | 20.8 ± 12.8 |
| Cooperative learning (no-PF) | 366.3 ± 152.4 | 1.6 ± 0.7 |
| Cooperative learning (ad-PF) | 334.9 ± 145.2 | 10.1 ± 12.1 |
| Robust Separate X | 478.3 ± 552.1 | 17.5 ± 10.6 |
| Robust Separate Z | 414.4 ± 423.4 | 14.6 ± 12.3 |
| Robust Early Fusion | 800.8 ± 1113.6 | 29.4 ± 15.5 |
| Robust Late Fusion | 504.6 ± 291.0 | 15.9 ± 12.7 |
| Robust Cooperative learning | **291.5 ± 149.3** | 50.8 ± 20.3 |

Robust Cooperative Learning also achieved a lower test MSE than both Cooperative Learning (no-PF) and Adaptive Cooperative Learning (ad-PF). In contrast, the other robust methods did not consistently improve predictive performance compared with their corresponding non-robust methods.

The number of selected features also varied substantially across methods. Robust Cooperative Learning tended to select more features than the other methods. Thus, in this real-data analysis, the advantage of the proposed method was observed primarily in terms of predictive performance rather than model sparsity.

## 6 Discussion and Conclusion

The experimental results demonstrate that incorporating robustness into the cooperative learning framework leads to improved predictive performance. A key reason for this improvement is that the proposed model explicitly accounts for deviations from the Gaussian noise assumption through the outlier absorption vector. In contrast, standard cooperative learning relies on a squared loss, which is sensitive to extreme observations[23]. As a result, even a small number of outliers can substantially affect the estimation of regression coefficients.

Another important observation is that the proposed method preserves the fundamental structure of cooperative learning. Specifically, the agreement parameter *ρ* continues to control the balance between early fusion and late fusion. This indicates that robustness can be incorporated without sacrificing the flexibility and interpretability of the original framework.

Furthermore, the results suggest that the proposed approach is particularly effective in high-dimensional settings with limited sample sizes. In such scenarios, combining sparsity-inducing regularization with robust estimation helps to stabilize the model and avoid overfitting.

Finally, the consistency between the simulation results and the real data analysis further supports the effectiveness of the proposed method.

## A Additional simulation results

### A.1 Predictive performance

**Figure 2:**
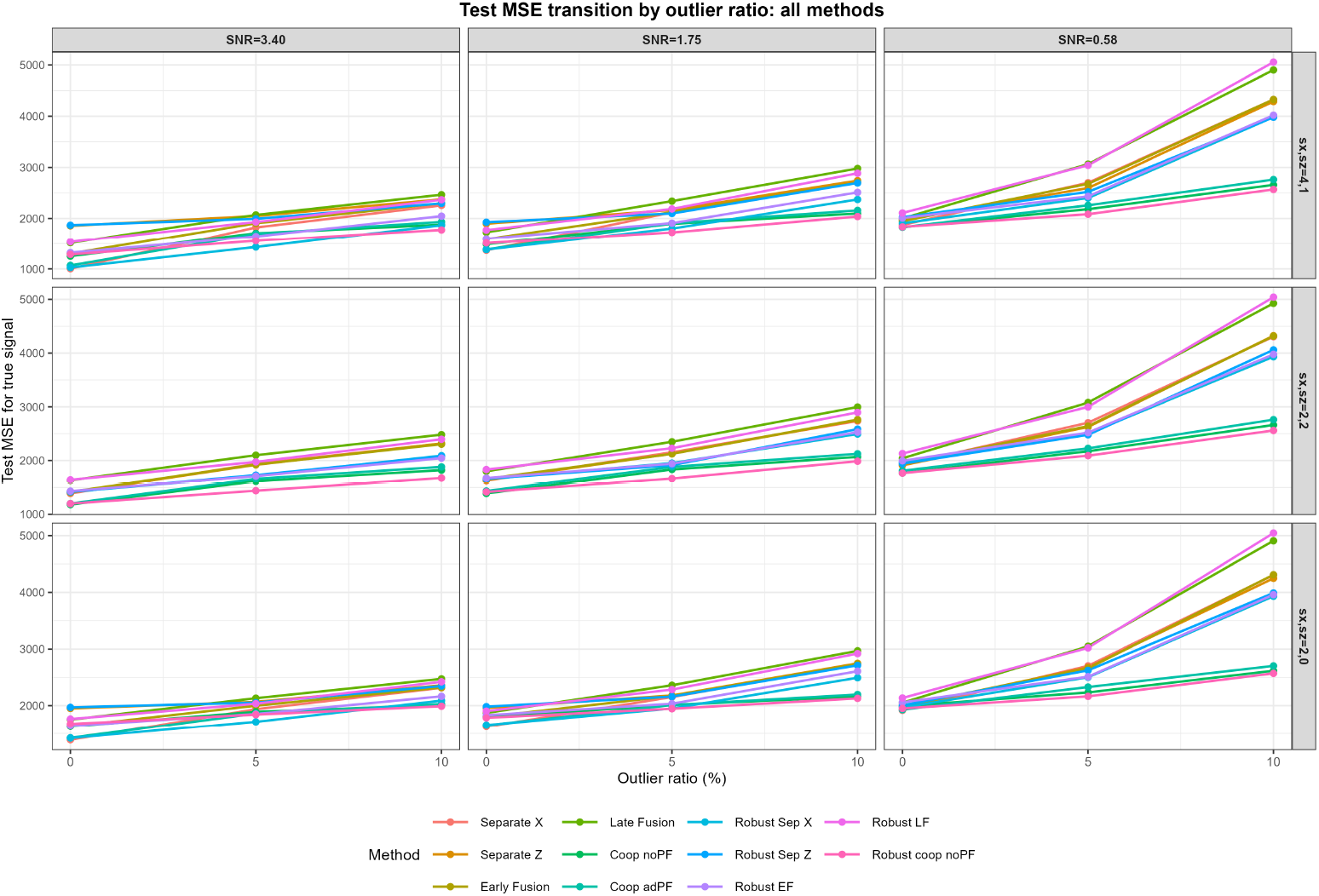
MSE transition for all methods.

### A.2 umber of selected features

**Figure 3:**
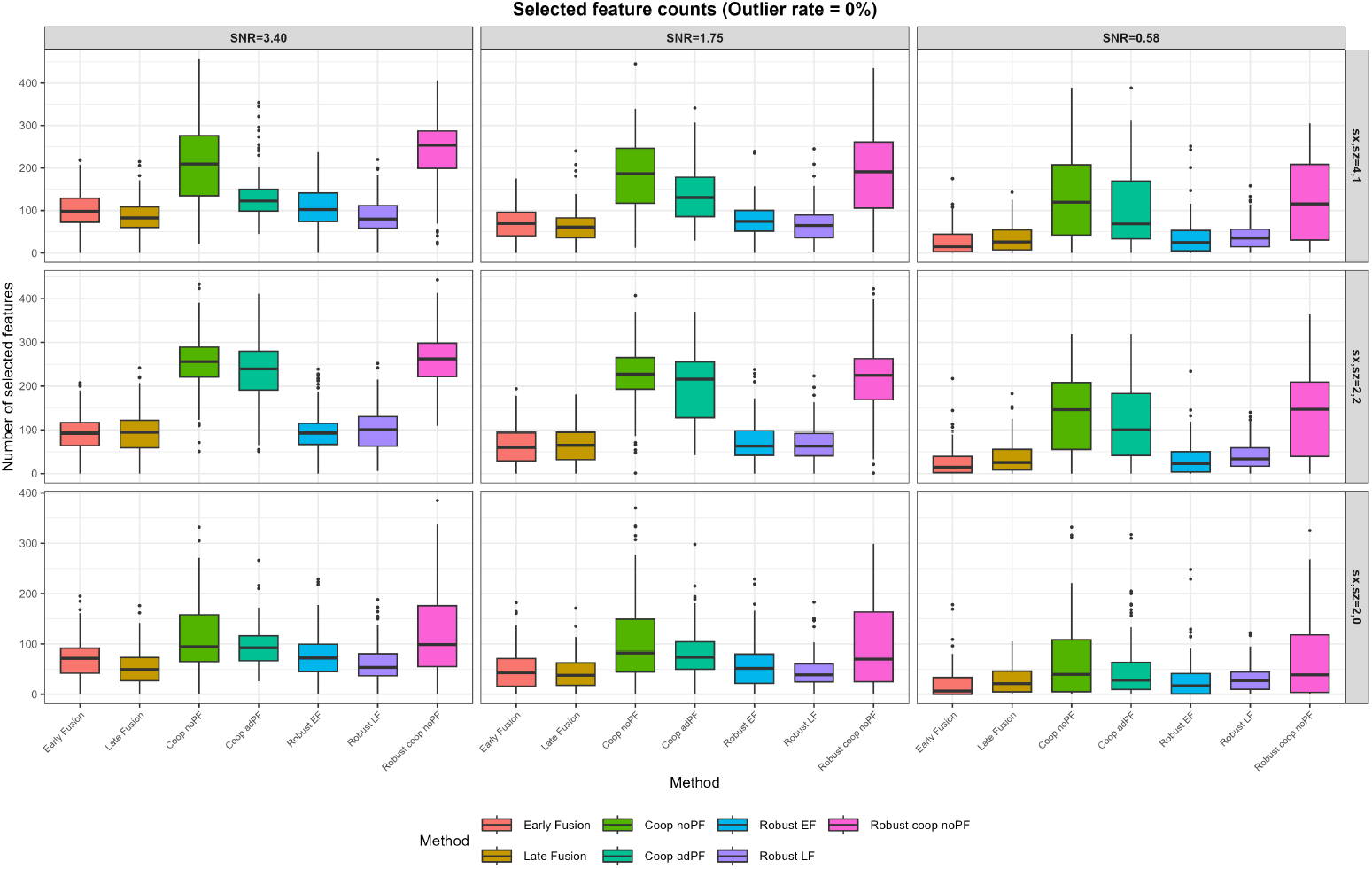
Numbers of Features outlier = 0%.

**Figure 4:**
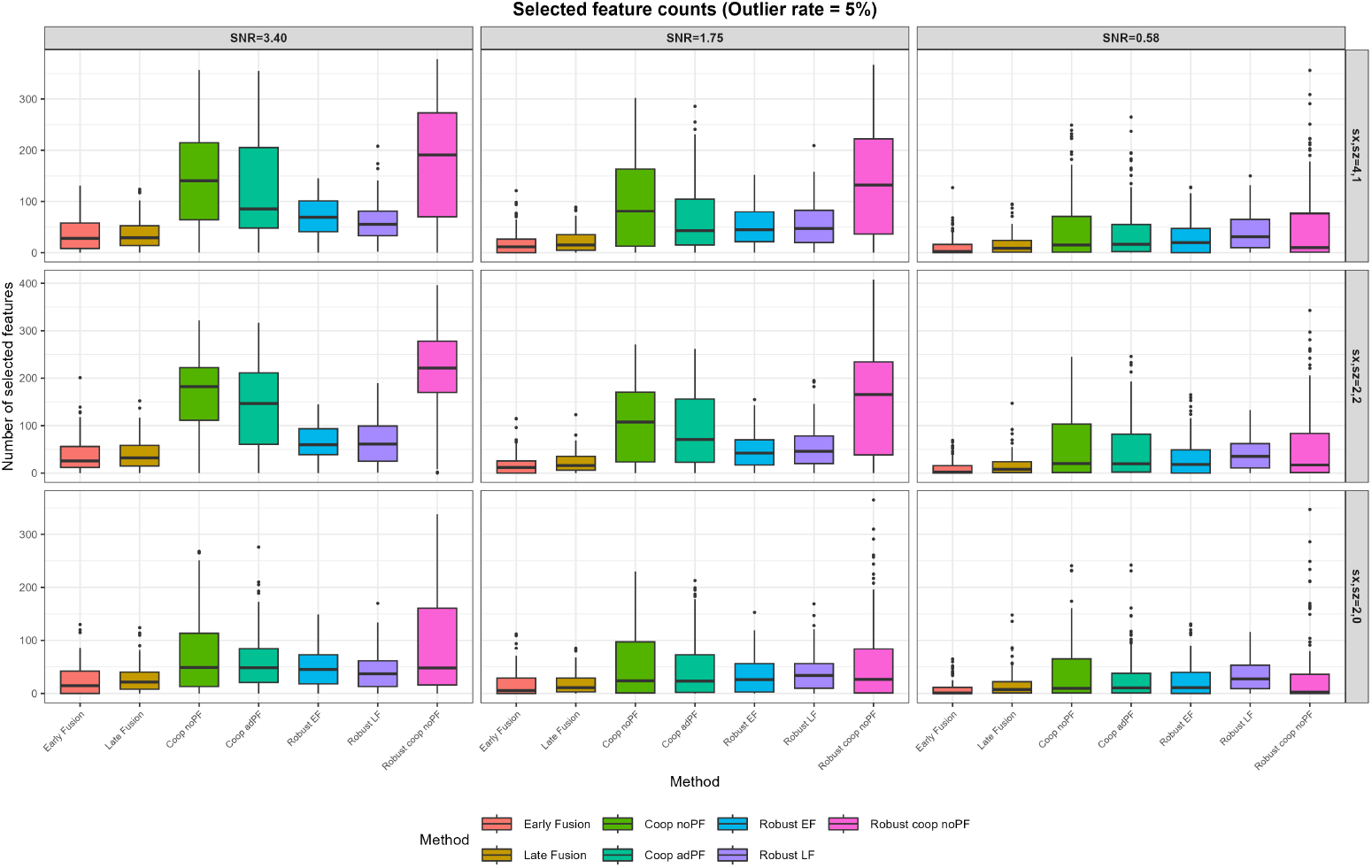
Numbers of Features outlier = 5%.

**Figure 5:**
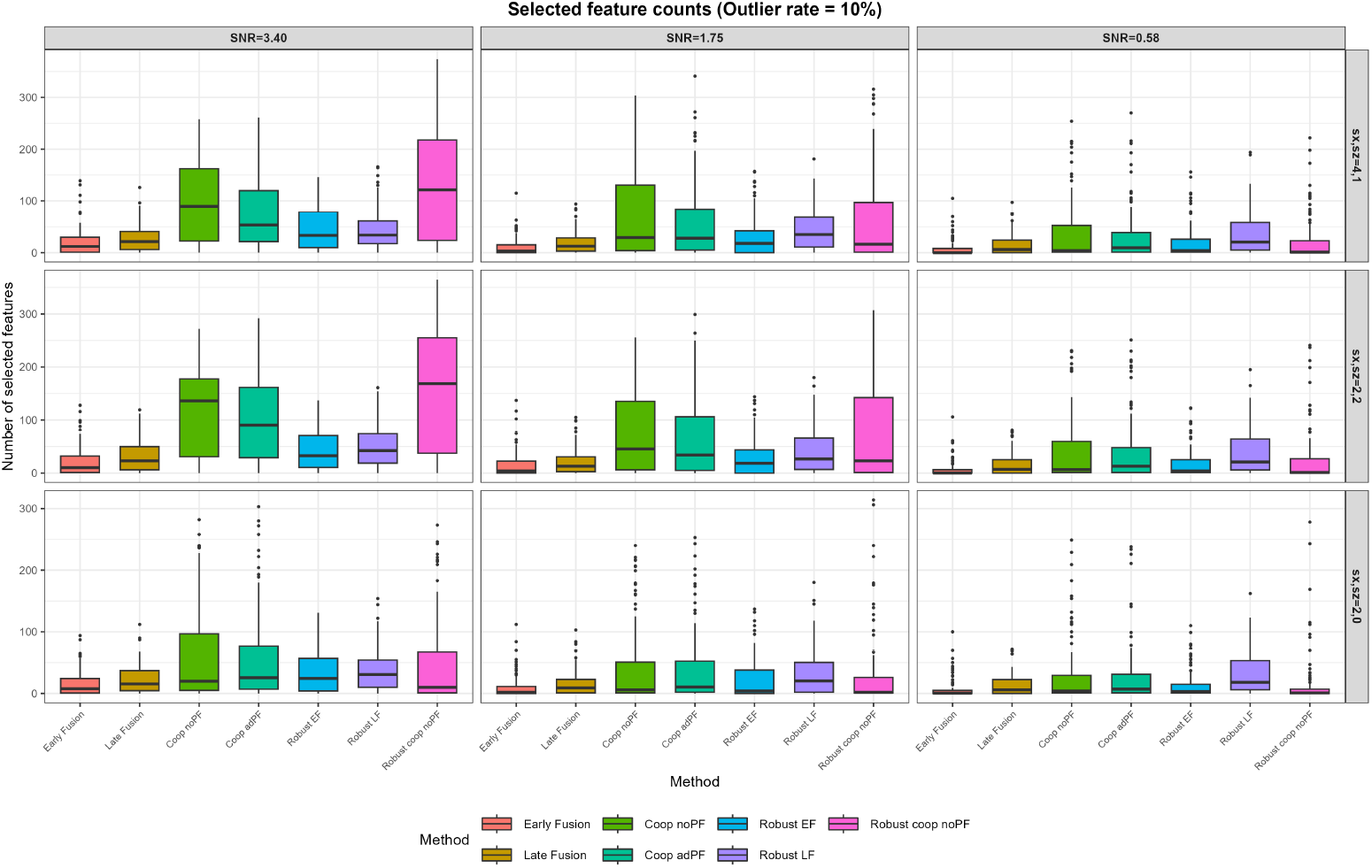
Numbers of Features outlier = 10%.

### A.3 Selection of the agreement parameter *ρ*

**Figure 6:**
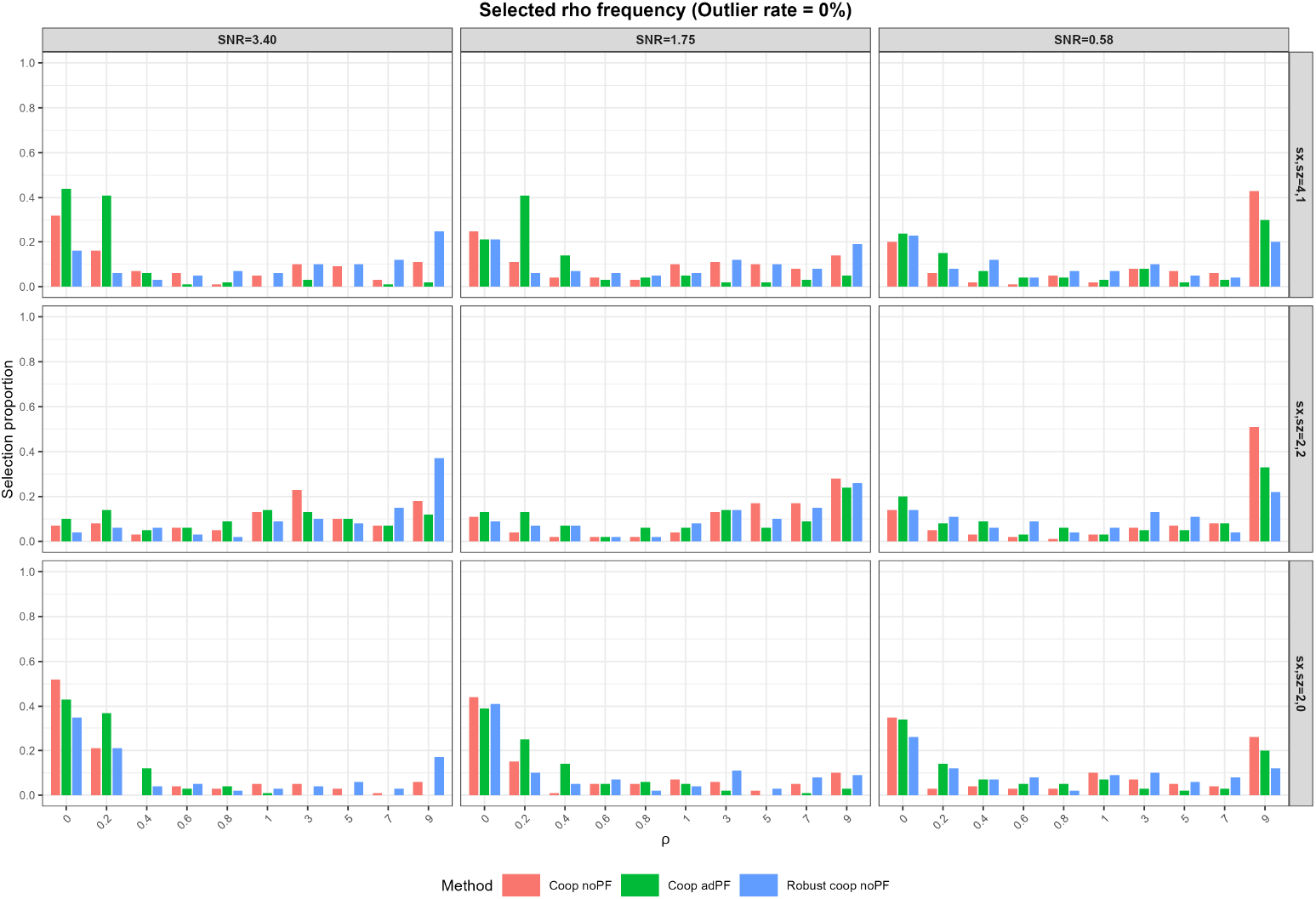
selected *ρ* outlier = 0%

**Figure 7:**
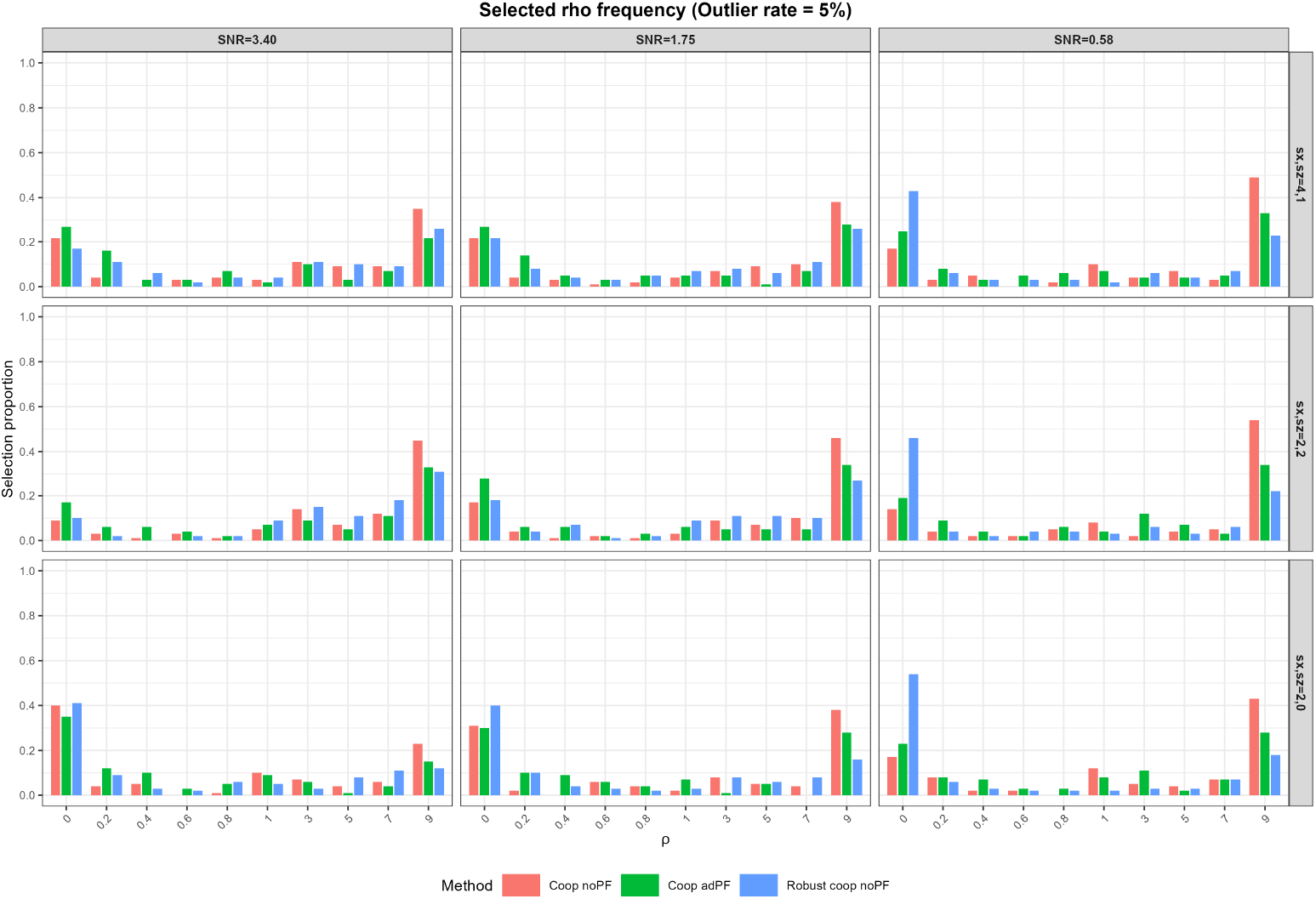
selected *ρ* outlier = 5%

**Figure 8:**
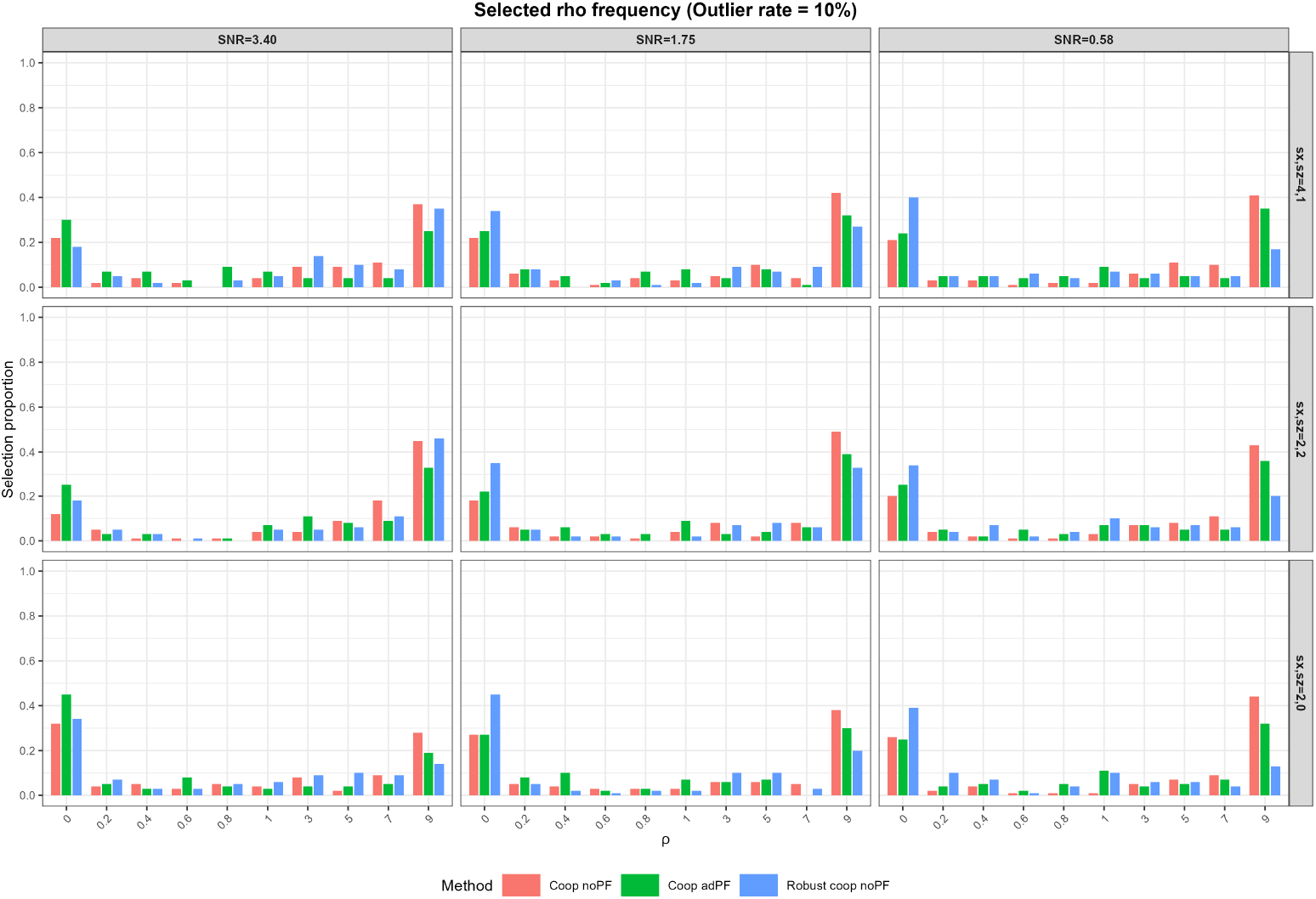
selected *ρ* outlier = 10%

